# [^3^H]5-MOP: a novel and selective colony stimulating factor-1 receptor (CSF1R) radiotracer

**DOI:** 10.64898/2026.05.12.724549

**Authors:** Camilla Iavazzo, Burcu A. Pazarlar, Benny Bang-Andersen, Thomas Jensen, Morten Hentzer, Jesper F. Bastlund, Kate Lykke Lambertsen, Bente Finsen, Anne M. Landau, Jens D. Mikkelsen

## Abstract

Colony stimulating factor 1 receptor (CSF1R) is a tyrosine kinase receptor that is expressed exclusively in microglia within the CNS. Its endogenous ligands, colony stimulating factor-1 (CSF1) and interleukin-34 (IL-34), are released from neurons, positioning CSF1R as a key mediator receptor of neuron-glia communication. CSF1R is considered not only a potential drug target, but also a biomarker of neuroinflammation. From that perspective, selective radioligands for neuroimaging are of great interest for imaging neuroinflammation and determining drug occupancy.

In this study, we have validated the binding characteristics of a CSF1R inhibitor, 4-((5-MethOxy-6-((5-methoxypyridin-2-yl)methoxy)pyridin-3-yl)methyl)-2-(1-methyl-1H-pyrazol-4-yl)pyrimidine (5-MOP) as a novel CSF1R radioligand, by performing *in vitro* saturation binding experiments in human and murine tissues. 5-MOP was found to be selective for CSF1R among a broad range of kinases. Autoradiography revealed that [^3^H]5-MOP binds with high affinity (K_D_ = 9.8 nM) to a single saturable binding site in human meningioma tissues, and this binding was displaced with known CSF1R inhibitors, including CPPC, sCSF1_inh_ and GW-2580. In contrast, CPPC, which has been extensively used as a CSF1R radioligand showed substantial cross-reactivity to other brain kinases, including Trk A/B/C, and [^3^H]CPPC could only be displaced with CPPC itself, not by other ligands, including 5-MOP. These results identify [^3^H]5-MOP as the most selective radioligand currently available, enabling accurate detection of drug occupancy and activated microglia.

**Significance of the study:** This study identifies and validates a novel selective radioligand that binds CSF1R with high selectivity and low nanomolar affinity. Because CSF1R is selectively expressed in activated microglia, this radioligand could be useful for detecting neuroinflammatory activity.

## Introduction

Colony stimulating factor 1 receptor (CSF1R, also known as M-CSFR, c-FMS, or CD-115) is a class III tyrosine kinase receptor (Chitu *et al*., 2021;Segaliny *et al*., 2015). It is activated by two distinct endogenous ligands: CSF1 (also known as macrophage-colony stimulating factor, M-CSF) and interleukin-34 (IL-34) (Lin *et al*., 2008;Wang *et al*., 2012) and they have distinct locations, and like mediate different functions to the receptor, and different effects in the microenvironment (Kana *et al*., 2019;Nandi *et al*., 2012). Conditional deletion of *Csf1r* in the brain combined with IL-34 knock-out in mice resulted in complete microglia deficiency (Kana *et al*., 2019). Macrophage lineage cells including microglia are the primary, if not only, cell type expressing CSF1R in the CNS as evidenced by in situ hybridization and promotor derived GFP expression in mice (Grabert *et al*., 2020;Sasmono *et al*., 2003).

CSF1R is a key regulator of microglia proliferation, differentiation, and survival (Jurga *et al*., 2020). The CSF1R kinase inhibitors have therefore been widely used as tools to deplete microglia (Green *et al*., 2020). Loss of microglia is also seen in mice with CRF1R knock out or after chronic pharmacological blockade (Elmore *et al*., 2014;Erblich *et al*., 2011) but these animals appeared to have a normal phenotype.

CSF1R has also been suggested as a drug target for multiple neurological disorders (El-Gamal *et al*., 2018). For example, autosomal dominant neurodegenerative disorder, such as adult-onset leukoencephalopathy with axonal spheroids and pigmented glia can be caused by CSF1R mutations (Du *et al*., 2025).

Upregulation of CSF1R is considered an indicator of activated microglia and observed in tissues from patients with Alzheimer’s disease (AD), multiple sclerosis, and glioblastoma (Akiyama *et al*., 1994). Single-cell transcriptomic analyses of several preclinical murine models of neuroinflammation demonstrate that CSF1 is upregulated in disease-associated microglia (Keren-Shaul *et al*., 2017). This prompted discovery and preclinical testing of various CSF1R inhibitors (Green *et al*., 2020). The small-molecule inhibitor sCSF1R_inh_ block CSF1R phosphorylation, which attenuates the inflammatory responses in both in vitro and in vivo studies (Hagan et al. 2020). These results support the evaluation of CSF1R-targeting approaches as a therapy for neuroinflammatory diseases, and CSF1R selective radioligands may be important tools in determining occupancy of drug development candidates binding to CRF1R. Several preclinical models of neurodegeneration of multiple sclerosis, AD, ALS, and neuronal injury support the notion that inhibiting CSF1R is beneficial in altering neuroinflammation (Borjini *et al*., 2016;Dagher *et al*., 2015;Hagan *et al*., 2020;Nissen *et al*., 2018;Spangenberg *et al*., 2019).

The current study has the overall aim of identifying novel radioligands for neuroinflammation, and CSF1R is a good candidate target with its restricted expression in microglia. CRF1R expression is increased in neuroinflammation, both in animal models of neurodegeneration and in postmortem tissues from patients with neurodegenerative diseases. For example, in transgenic models of AD-like pathology CSF1R-dependent progressive increases specifically in microglial proliferation in the proximity to Aβ-plaques (Olmos-Alonso *et al*., 2016) highlighting the potential value of CSF1R imaging as a tool in studying pathology-induced neuroinflammation. Until now, translocator protein, 18kDa (TSPO) is the most widely used target for positron emission tomography (PET), but it is not optimal due to genetic variations, non-microglia expression, and lack of detecting certain changes in inflammation (Chauveau *et al*., 2021;Chauveau *et al*., 2025). In recent years, several potential CSF1R radioligands have been identified, including CPPC, GW-2580 and more recently JNJ-CSF1R-1 (Altomonte *et al*., 2023;Bernard-Gauthier and Schirrmacher, 2014;Horti *et al*., 2019;Lee *et al*., 2022;Ogata *et al*., 2022;Tanzey *et al*., 2018;van der Wildt *et al*., 2022;Zhou *et al*., 2021) with CPPC being the most extensively used (Adhikari *et al*., 2024;Chauveau *et al*., 2025). However, no CSF1R PET radioligand has been demonstrated to be completely specific for CSF1R or clinically effective for *in vivo* imaging of CSF1R in human brain.

In this perspective, there is still an unmet need for a validated and selective CSF1R radioligand. Here, we report here the discovery of a novel radioligand, [^3^H]5-MOP, based on a compound previously described in the patent literature (Richards, 2021) and validated its binding properties in comparison to CPPC in human and murine tissues. Owing to its selectivity, we further examined its utility in a mouse model of cerebral ischemia with extensive neuroinflammation.

## Materials and Methods

### Radioligands and compounds

[^3^H]-5-MOP was radiolabelled by RC Tritec (Teufen, Switzerland). The specific activity of the radioligand was 101.6 Ci/mmol. The radiochemical concentration was 1 mCi/mL on the day of synthesis. [^3^H]-CPPC was radiolabelled by Novandi Chemistry AB (Södertalje, Sweden). The specific activity of the radioligand was 83 Ci/mmol. The radiochemical concentration was 1 mCi/mL on the day of synthesis.

[^3^H]-PBR28 was radiolabelled by RC Tritec (Teufen, Switzerland). The specific activity of the radioligand was 80.7 Ci/mmol. The radiochemical concentration was 1 mCi/mL on the day of synthesis.

The compounds used for the displacement of [^3^H]-5-MOP and [^3^H]CPPC were 5-MOP ([4-((5-methoxy-6-((5-methoxypyridin-2-yl)methoxy)pyridin-3-yl)methyl)-2-(1-methyl-1H-pyrazol-4-yl)pyrimidine, batch A1628DD), CPPC (5-cyano-N-(4-(4-[11C]methylpiperazin-1 yl)-2-(piperidin-1-yl)phenyl)furan-2-carboxamide, batch A1628OL), sCSF1R_inh_ ((S)-4-(3-((2-(6-methoxypyridin-3-yl)-2,3-dihydrobenzo[b]dioxin-6-yl)methyl)-3H-imidazo[4,5-b]pyridin-6-yl)-2-methylbut-3-yn-2-amine, batch A1710WS), and GW-2580 (5-(3 Methoxy-4-((4-methoxybenzyl)oxy)benzyl)pyrimidine-2,4-diamine, batch A1627XW) and for [^3^H]PBR28 (N-([3H]2-methoxybenzyl)-N-(4-phenoxypyridin-3-yl)acetamide) it was PK11195 (1-(2-chlorophenyl)-N-methyl-N-(1-methylpropyl-3-isoquinoline carboxamide, batch A0031BF) all synthesised by H. Lundbeck A/S (Valby, Denmark).

### CSF1R Kinase Binding assay

CSF1R kinase binding was determined using LanthaScreen Kinase Binding assay from ThermoFisher Scientific (Table 1) The assay was set up using the following reagents: Kinase: wildtype CSF1R (BPS Bioscience, cat.no. 40227);kinase tracer 236, (ThermoFisher Scientific;cat.no. PV5592);LanthaScreen™Eu-anti-His Antibody, (ThermoFisher Scientific cat.no. PV5597).

**Table 1.**
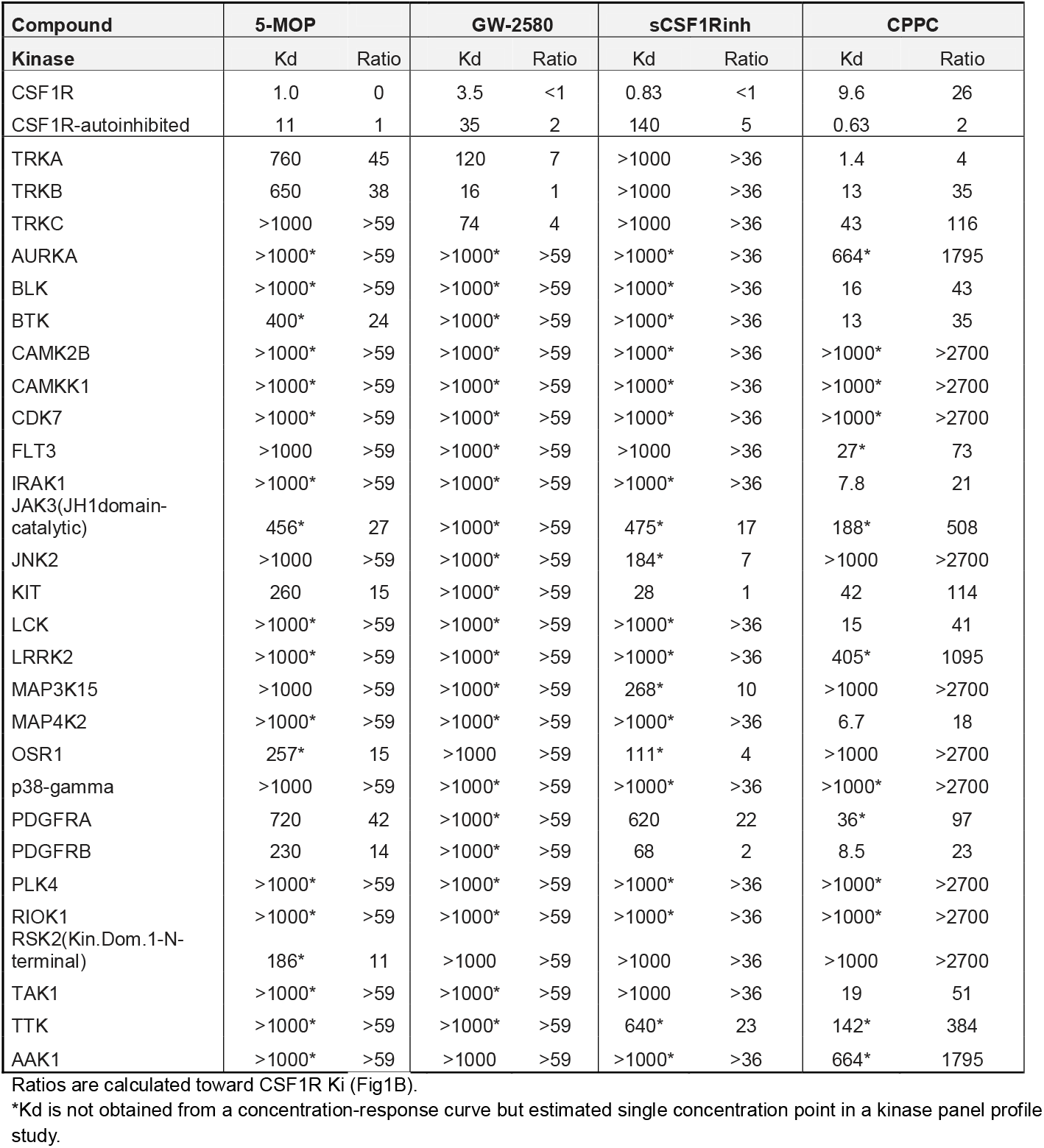
Broad kinase binding profile and kinase selectivity for 5-MOP and key literature compounds. Ratio is calculated relative to the CSF1R K_i_ value from Fig. 1B. Of note, K_d_ values are not obtained from a concentration-response curve, but estimated from a single concentration point in a kinase panel profile study. The K_i_ value shows the best correlation to cellular IC_50_ values rather than the CSF1R non-autoinhibited and autoinhibited forms which represent different truncations and extreme conformations of CSF1R (see methods).

**Fig. 6**
| Compound | 5-MOP |  | GW-2580 |  | sCSF1Rinh |  | CPPC |  |
| --- | --- | --- | --- | --- | --- | --- | --- | --- |
| Kinase | Kd | Ratio | Kd | Ratio | Kd | Ratio | Kd | Ratio |
| CSF1R | 1.0 | 0 | 3.5 | <1 | 0.83 | <1 | 9.6 | 26 |
| CSF1R-autoinhibited | 11 | 1 | 35 | 2 | 140 | 5 | 0.63 | 2 |
| TRKA | 760 | 45 | 120 | 7 | >1000 | >36 | 1.4 | 4 |
| TRKB | 650 | 38 | 16 | 1 | >1000 | >36 | 13 | 35 |
| TRKC | >1000 | >59 | 74 | 4 | >1000 | >36 | 43 | 116 |
| AURKA | >1000* | >59 | >1000* | >59 | >1000* | >36 | 664* | 1795 |
| BLK | >1000* | >59 | >1000* | >59 | >1000* | >36 | 16 | 43 |
| BTK | 400* | 24 | >1000* | >59 | >1000* | >36 | 13 | 35 |
| CAMK2B | >1000* | >59 | >1000* | >59 | >1000* | >36 | >1000* | >2700 |
| CAMKK1 | >1000* | >59 | >1000* | >59 | >1000* | >36 | >1000* | >2700 |
| CDK7 | >1000* | >59 | >1000* | >59 | >1000* | >36 | >1000* | >2700 |
| FLT3 | >1000 | >59 | >1000* | >59 | >1000 | >36 | 27* | 73 |
| IRAK1 | >1000* | >59 | >1000* | >59 | >1000* | >36 | 7.8 | 21 |
| JAK3(JH1domain-catalytic) | 456* | 27 | >1000* | >59 | 475* | 17 | 188* | 508 |
| JNK2 | >1000 | >59 | >1000* | >59 | 184* | 7 | >1000 | >2700 |
| KIT | 260 | 15 | >1000* | >59 | 28 | 1 | 42 | 114 |
| LCK | >1000* | >59 | >1000* | >59 | >1000* | >36 | 15 | 41 |
| LRRK2 | >1000* | >59 | >1000* | >59 | >1000* | >36 | 405* | 1095 |
| MAP3K15 | >1000 | >59 | >1000* | >59 | 268* | 10 | >1000 | >2700 |
| MAP4K2 | >1000* | >59 | >1000* | >59 | >1000* | >36 | 6.7 | 18 |
| OSR1 | 257* | 15 | >1000 | >59 | 111* | 4 | >1000 | >2700 |
| p38-gamma | >1000 | >59 | >1000* | >59 | >1000* | >36 | >1000* | >2700 |
| PDGFRA | 720 | 42 | >1000* | >59 | 620 | 22 | 36* | 97 |
| PDGFRB | 230 | 14 | >1000* | >59 | 68 | 2 | 8.5 | 23 |
| PLK4 | >1000* | >59 | >1000* | >59 | >1000* | >36 | >1000* | >2700 |
| RIOK1 | >1000* | >59 | >1000* | >59 | >1000* | >36 | >1000* | >2700 |
| RSK2(Kin.Dom.1-N-terminal) | 186* | 11 | >1000 | >59 | >1000 | >36 | >1000 | >2700 |
| TAK1 | >1000* | >59 | >1000* | >59 | >1000 | >36 | 19 | 51 |
| TTK | >1000* | >59 | >1000* | >59 | 640* | 23 | 142* | 384 |
| AAK1 | >1000* | >59 | >1000 | >59 | >1000* | >36 | 664* | 1795 |

Ratios are calculated toward CSF1R Ki (Fig1B).
\*Kd is not obtained from a concentration-response curve but estimated single concentration point in a kinase panel profile study.

The assay is a homogeneous time-resolved fluorescence resonance energy transfer-based assay that measures binding of kinase tracer 236 to a GST/His-tagged CSF1R kinase. The resulting TR-FRET ratio correlates with kinase binding. Compounds that displace tracer binding will cause a decrease in TR-FRET ratio. The CSF1R kinase was obtained from BPS Bioscience and comprises residue 538 to 972 of the full-length human wildtype CSF1R kinase and contains a N-terminal GST-His-tag. The kinase binding reactions were performed in a 15 μL volume in 384-well plates. The kinase binding buffer A (5X Kinase Buffer A, cat.no. PV3189. ThermoFisher Scientific) consisted of 50 mM HEPES pH 7.5, 10 mM MgCl2, 1 mM EGTA, 0.01%Brij-35. In the assay, 1 or 2 nM CSF1R GST-tagged kinase in reaction buffer was incubated with the test compound (typically at 0 to 2000 nM), 30 or 60 nM tracer 236 and 0.2 nM detection Eu-labeled anti-GST antibody (included in kit) for 60 minutes at RT (15 μL total volume). TR-FRET 620/655 signal ratio was measured on a BMG Labtech Pherastar FSX multimode microplate reader with TRF laser for excitation. The signal integration time was 200 μs and start time 100 μs after excitation.

The TR-FRET ratio readout for test compounds was normalized to 0%inhibition corresponding to signal measured in control wells (median signal) with no inhibition of the kinase activity and 100%inhibition corresponding to signal measured in control wells with inhibitor (median signal). Test compound potency (K_i_,app) was estimated by nonlinear regression using the tight-binding fit model in GeneData Screener (GeneData AG). The tight-binding fit model is: SS=S0-(S_inf_-S0)*(([E]+[I]+K_i_,app-([E]+[I]+K_i_,app)2 - 4*[E]*[I])0.5) / (2*[E]))

Where S0 is the estimated efficacy (%inhibition) at infinite compound dilution, and S_inf_ is the maximal efficacy (%inhibition), [E] is the enzyme concentration (active enzyme concentration) and [I] is the concentration of test compound. K_i_,app is the apparent dissociation constant of the inhibitor.

### Broad Kinase Binding assay

The kinase selectivity was evaluated using the KINOMEscan™active site-directed competition-binding methodology (Eurofins/DiscoveRx) (Fabian et al. 2005). This method reports thermodynamic interaction affinities between test compounds and kinases in the absence of ATP. Compounds that bind the kinase active site (and prevent kinase binding to an immobilized ligand) reduce the amount of kinase captured on a solid support. Screening hits are identified by measuring the amount of kinase captured in test versus control samples by using a quantitative polymerase chain reaction (PCR) method that detects an associated DNA label. The CSF1R, autoinhibited form is comprised of residues 538-939, including the JM-B, JM-S and JM-Z domains, whereas the CSF1R, non-autoinhibited form is comprised of residues 564-939, including the JM-Z domain upstream of the kinase domain (Wodicka et al. 2010).

### Tissues used in for binding studies

Tissue samples obtained from four (2 females;2 males) human supratentorial intracranial meningioma patients that underwent neurosurgical resection in the period of September to October 2023 were used in this investigation. Serial cryostat sections from the four tumour resections were incubated with either [^3^H]5-MOP or [^3^H]PBR28 used as radiotracer for translocator protein 18kDa (TSPO), a marker of microglia, at concentration 24 and 3 nM, respectively. Based on this, sections from the sample with most intense labelling were used for further validation experiments as described below.

Induction of experimental ischemic stroke was performed in 8-week-old, male C57BL/6 mice (Janvier Labs, Le Genest-Saint-Ilse, France). Animals were housed at the Biomedical Laboratory, University of Southern Denmark and allowed to acclimatize at least one week prior to experiments. Mice were group-housed (3-4 mice/cage) under controlled temperature and humidity, with *ad libitum* access to food and water and a 12-hour light/dark cycle. All procedures were approved by the Danish Animal Inspectorate (J. no 2023-15-0201-01603). Permanent middle cerebral artery occlusion (pMCAO) was performed as previously described (Lambertsen *et al*., 2001). Briefly, mice were anesthetized using Hypnorm (fentanyl citrate 0.315 mg/ml and fluanisone 10 mg/ml, VetaPharma Ltd, Leeds, UK), Midazolam (5 mg/ml, Hameln Pharma, Hameln, Germany), and distilled water (1:1:2). Following incision and dissection of the temporal muscle, the left MCA was exposed and permanently occluded by microbipolar coagulation. Mice were sutured, given s.c. 0.9%saline to prevent dehydration, and maintained in a 28_C_ heating cabinet for 24 hours. Post-operative analgesia consisted of s.c. buprenorphine hydrochloride (0.001 mg/20 g, Temgesic) administered three times at eight-hour interval starting at surgery. Mice were euthanized by cervical dislocation at day 5 after pMCAO (n=8), when the ischemic core is surrounded by activated microglia (Ma *et al*., 2020). Brains were rapidly removed, frozen in gaseous CO_2_, and cryosectioned as described for human samples.

Cryostat sections (16 µm) were cut at -20°C using a Epredia CryoStarTM NX70 cryostat (Thermo ScientificTM, Waltham, USA) and mounted on pre-coated glass slides (Superfrost™ Plus Adhesion Microscope Slides). The tissue slices were thaw-mounted on each glass slide (SuperfrostTM Plus Adhesion Microscope Slides, Thermo Fisher Scientific, Waltham, USA). The slides were stored at -20°C until the start of the experiment.

### Autoradiography for [^3^H]5-MOP and [^3^H]-CPPC. Saturation and inhibition studies

Glass slides with two frozen adjacent tissue sections from the same sample were allowed to reach room temperature and then preincubated twice for 10 minutes in 50 mM Tris-HCl buffer (pH 7.4), containing 0.5%bovine serum albumin (BSA). After the preincubation step, the incubation buffer with the radioligand was gently pipetted onto the glass slides, which were placed on a rotator at room temperature for 120 minutes. For the saturation study, increasing concentrations of the radioligand [^3^H]5-MOP (0.31 nM, 0.62 nM, 1.25 nM, 2.5 nM, 5 nM, 10 nM, 20 nM, 30 nM) were added to the incubation buffer, which contains 50 mM Tris-HCl (pH 7.4), 5 mM MgCl_2_, 2 mM EGTA and 0.5%BSA. For [^3^H]-CPPC increasing concentrations (0.31 nM, 0.62 nM, 1.25 nM, 2.5 nM, 5 nM, 10 nM, 20 nM, 30 nM) were dissolved in the same buffer. Non-specific binding was determined by adding either 10 µM 5-MOP or 10 µM CPPC to the incubation solution.

For the inhibitory binding study, increasing concentrations of three CSF1R competitors (5-MOP, CPPC, sCSF1R_inh_, and GW-2580) were tested at 0 nM, 0.5 nM, 1 nM, 2 nM, 4 nM, 10 nM, 20 nM, 40 nM, 100 nM, 500 nM, 2,000 nM, and 10,000 nM. These were added to the incubation buffer, in the presence of 24 nM of [^3^H]5-MOP radioligand.

Sections from mice were processed as described above using either 3 nM [^3^H]PBR28, 24 nM [^3^H]-CPPC, or 24 nM [^3^H]-5MOP.

After slides were incubated in an incubation buffer containing radioligand, then they were washed three times in an ice-cold preincubation buffer for 5 minutes each, followed by a final rinse in 4°C distilled water for 10 seconds. The slides were then air-dried for 45 minutes in a fume hood with normal air circulation. Subsequently, slides were dried overnight in a paraformaldehyde chamber and exposed to FUJI imaging phosphor plates for 72 hours at 4°C, alongside with different [^3^H] standards. The reference standards used were [^3^H] microscale ART0123 (0−489.1 nCi/mg), ART0123B (3−109.4 nCi/mg), ART0123C (0.10-15.9 nCi/mg), and purchased from American Radiolabeled Chemicals, Inc., St. Louis, USA), while [^3^H] microscale Batch 21A RPA510 was purchased from GE Healthcare, UK. The FUJI imaging plates were scanned by Fujifilm Image Reader (BAS-2500 V1.8), and the autoradiograms were imaged and analyzed.

### Data analysis

Quantitative receptor binding analysis was conducted using ImageJ software (version 1.53t, U. S. National Institutes of Health, Bethesda, USA), The entire section was delineated, and measurements were taken from both sections from each glass slide. The calculated grey value optical densities were converted to standards of known concentrations (nCi/mg), and interpolated values were converted into the amount of bound radioligand (fmol/mg of Tissue Equivalent, TE) and the mean of the technical replicates was then calculated for each data point. Specific binding was calculated as the difference between total binding and the non-specific binding. The K_D_, B_max_, and K_i_ values were calculated using GraphPad Prism (version 9.5.1, GraphPad Software, San Diego, USA).

## Results

### Biochemical characterisation of 5-MOP

The chemical structures of 5-MOP and the other CSF1R inhibitors investigated in the present study are shown in Fig. 1A. As illustrated, 5-MOP shares structural features with GW-2580, whereas it is structurally distinct from that of CPPC, sCSF1R_inh_, and JNJ-CSF1R-1.

**Figure 1.**
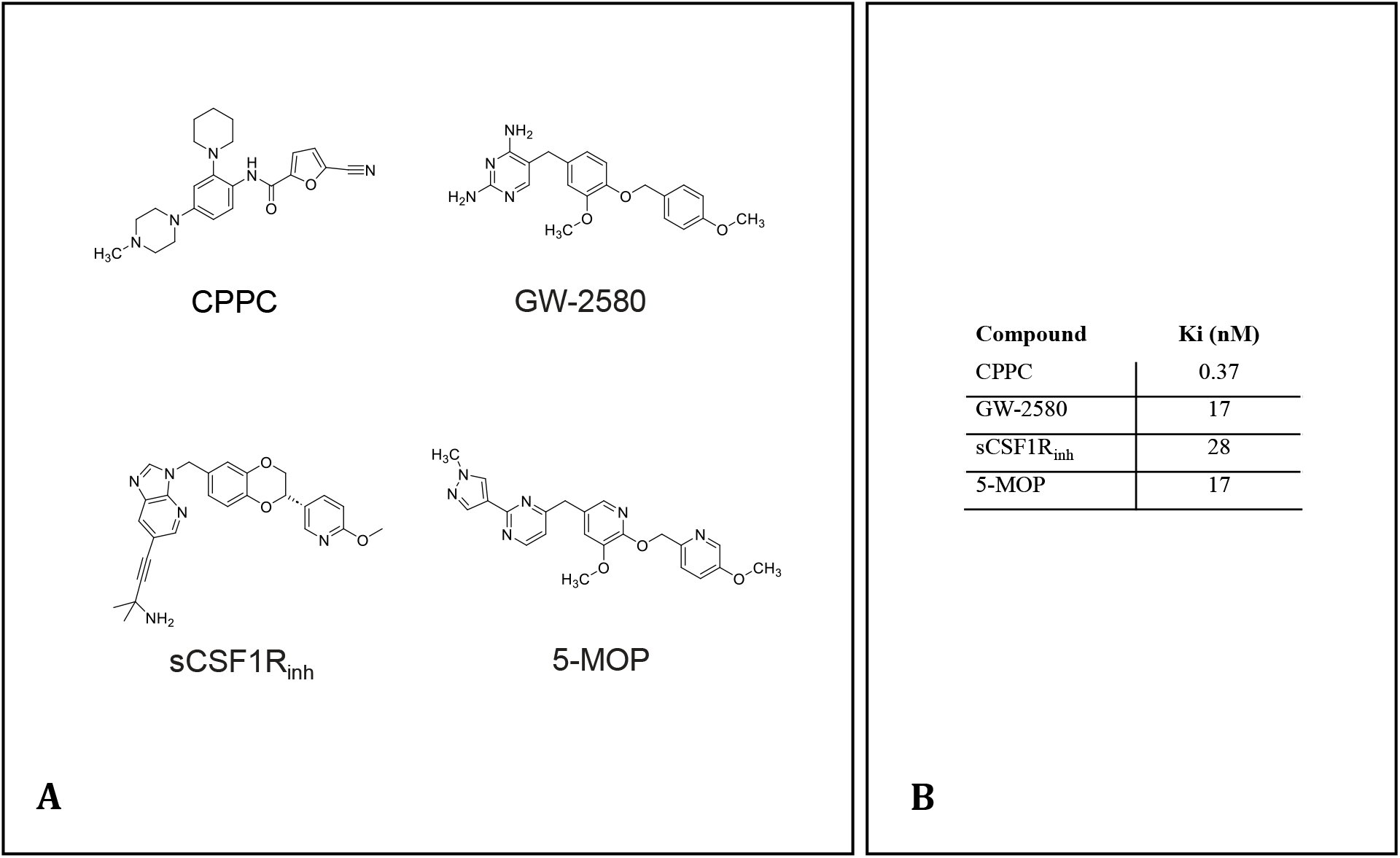
Structure and activity of CSF1R modulators. Molecular structure of CPPC, GW-2580, sCSF1R_inh_, and 5-MOP (A), as well as their K_i_ in inhibition of CSF1R activity (B).

The compounds were evaluated in a CSF1R LanthaScreen kinase binding ssay (Fig 1B) and against a panel of kinases using the KINOMEscan™ binding assay (Table 1). The potency in the LanthaScreen kinase binding assay is reported as K_i_ values (nM), whereas it is reported as K_d_ values (nM) in the and estimated for each compound in both the autoinhibited and non-autoinhibited states. The ratios shown in Table 1 is the calculated kinase K_d_ value relative to the CSF1R K_i_ value shown in Fig. 1B.

All four compounds exhibited low-nanomolar potency (K_i_ values 17-28 nM) in the LanthaScreen kinase binding assay, whereas CPPC demonstrated sub-nanmolar potency (K_i_ = 0,37 nM). In the KINOMEscan™ binding assay, all four compounds were tested against a panel of 406 different kinases. Only 5-MOP showed no off-target binding defined as binding ratio relative relative to CSF1R K_i_. Notably, 5-MOP showed no measurable activity at any other tested kinases tested at concentrations below 200 nM. In contrast, the other three compounds exhibited activity at additional kinases, with potencies comparable to their activity at CSF1R. In particular, CPPC and GW-2580 showed substantial activity Trk receptors that are highly expressed in the CNS.

### Binding of [^3^H]5-MOP to human tissues is saturable and selective at low nM concentrations

[^3^H]5-MOP was investigated for its binding characteristics in frozen, non-fixed tissue sections obtained from neurosurgically resected human meningioma. In an initial screening step, [^3^H]PBR28, a selective radioligand for TSPO commonly used as a microglial marker, was used to identify the specimens with the highest binding, based on the observed positive correlation between the total binding of [^3^H]-PBR28 and [^3^H]5-MOP. As shown in Fig. 2A, [^3^H]PBR28 displayed robust binding to the tumour tissue, which was almost completely displaced in the presence of PK11195, another TSPO ligand (Fig. 2A). The samples exhibiting the strongest [^3^H]PBR28 signal were subsequently selected for further characterisation with [^3^H]5-MOP.

**Figure 2.**
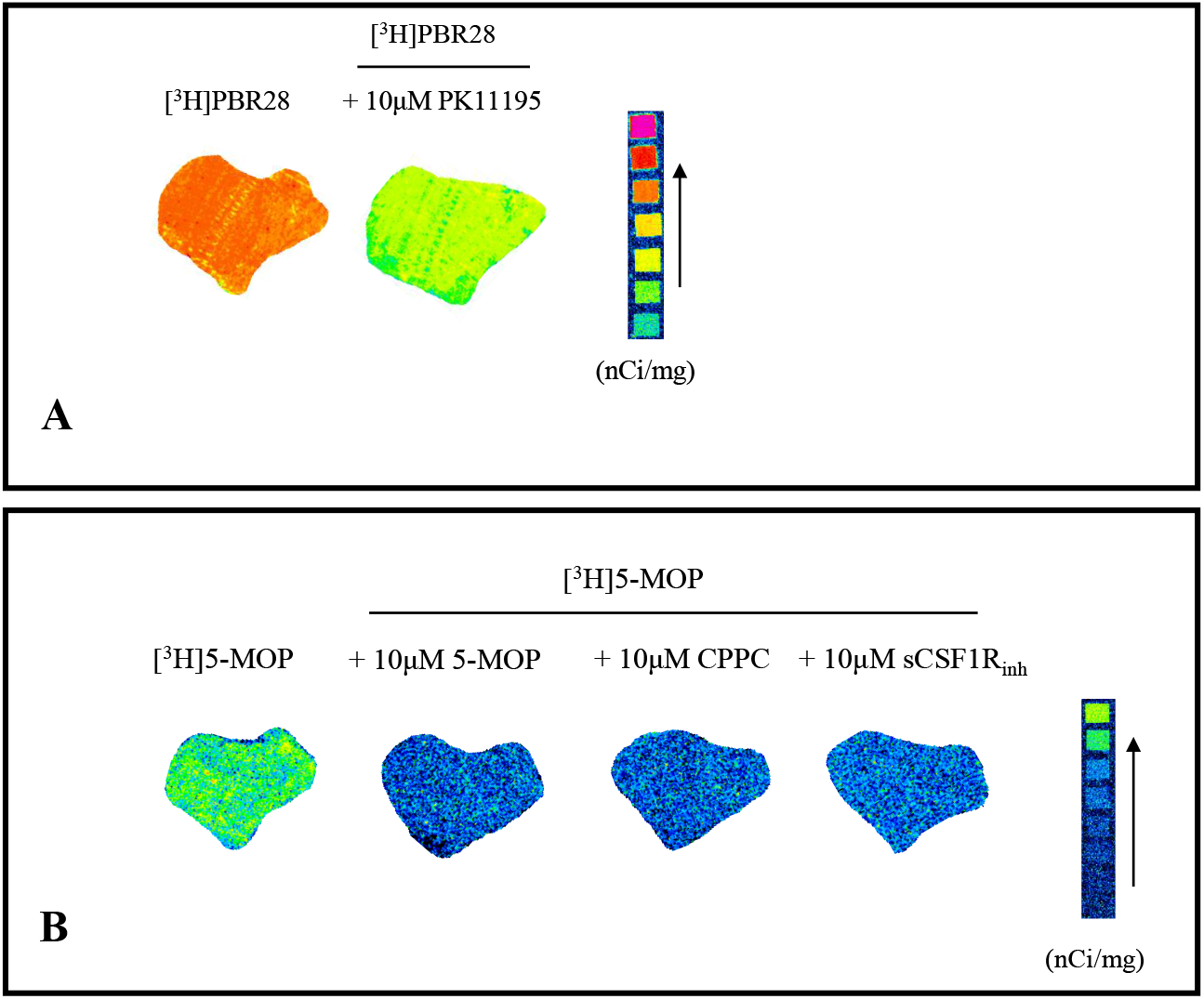
Representative autoradiograms of [^3^H]5-MOP and [^3^H]-PBR28 binding to human meningioma tissue. Sections from one meningioma resection were used to assess the binding to CSF1R and TSPO. Figure 2A shows autoradiographic images for [^3^H]-PBR28 with and without displacement with another TSPO ligand, PK11195 Figure 2B shows autoradiograms from the same resection with [^3^H]5-MOP alone and after displacement with 10 µM 5-MOP, CPPC, or sCSF1R_inh_. Notably, [^3^H]5-MOP bound to the tissue section, and all cold molecules were able to completely block the binding. Radioligand binding densities (nCi/mg) are shown below sections, with a standard for qualitative comparison.

[^3^H]5-MOP binding at various conditions was evaluated on adjacent sections from the same sample tissue (Fig. 3). First, saturation binding of [^3^H]5-MOP was performed on two selected tissue samples, both in the absence and presence of 5-MOP as well as the three other CRF1R ligands that were characterised biochemically. The saturation binding curves of the [^3^H]5-MOP shown in Fig. 3 all demonstrated a profile consistent with a saturable binding site. Displacement with high concentration of un-labelled 5-MOP produced, as expected, full displacement of [^3^H]5-MOP, and the resulting non-specific binding curve was linear (Fig. 3A). The calculated K_D_ value was 9.8 nM.

**Figure 3.**
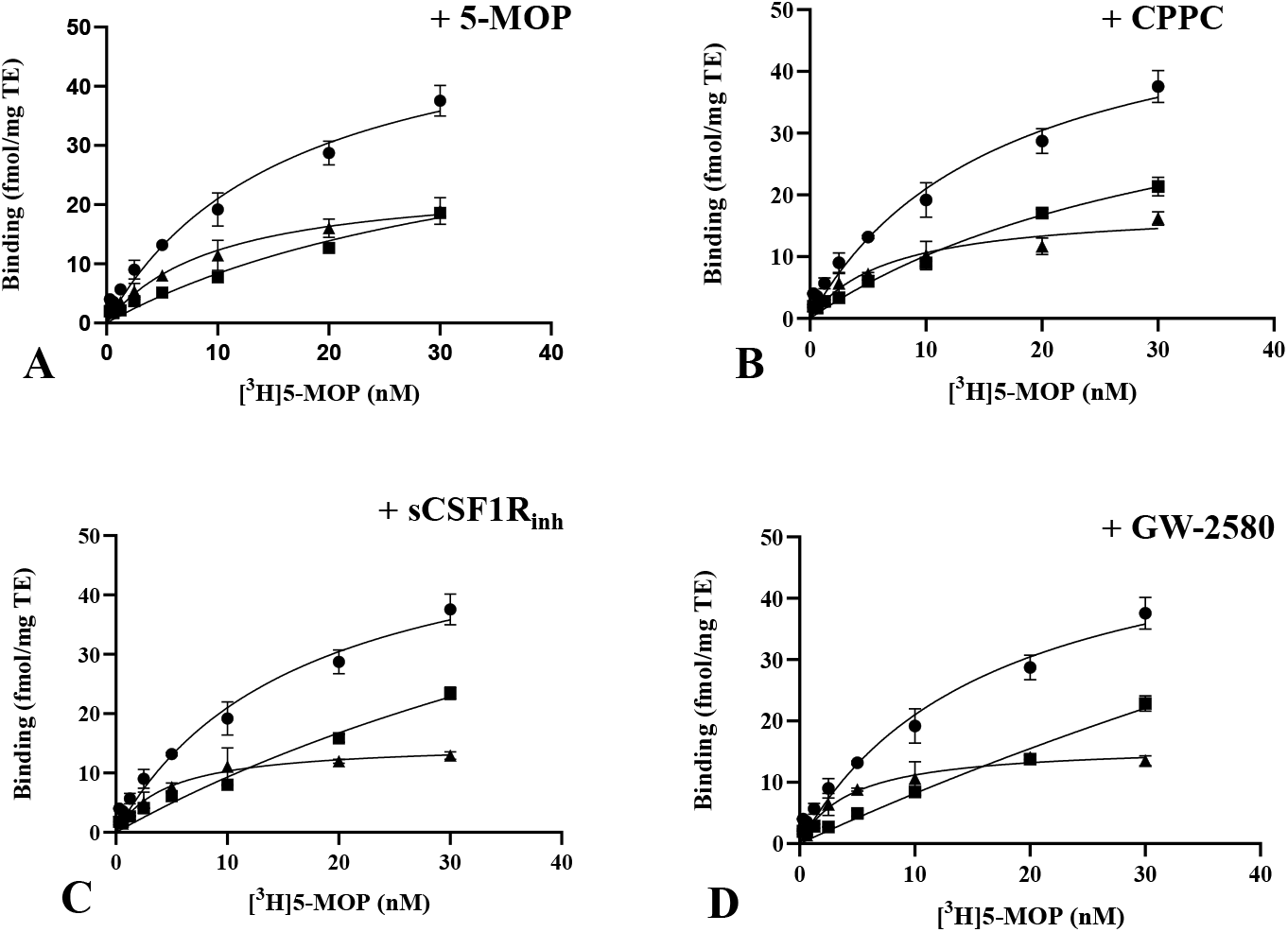
Saturation binding of [^3^H]5-MOP. Saturation binding curves for [^3^H]5-MOP (concentration range from 0,31 to 30 nM) in human meningioma tissue are shown under controlled conditions and in the presence of 10 µM 5-MOP (A), 10 µM CPPC (B), 10 µM sCSF1R_inh_ (C), or 10 µM GW-2580 (D). In all panels, the addition of the competing ligand displaced total binding, allowing separation of the specific and non-specific components. Specific binding curves show ligand saturation K_D_ values in low nanomolar range. For all the graphs, the x-axis represents radioligand concentrations (nM), and the y-axis represents binding (fmol/mg Tissue Equivalent, TE). ⍰= total binding;⍰= specific binding;⍰= non-specific binding.

While displacement by the un-labelled compound itself perhaps is expected, specificity is more convincingly demonstrated when structurally different ligands are also able to displace the radiolabeled molecule. CPPC was applied at high concentrations, and [^3^H]5-MOP was displaced with a K_D_ estimated to be 6.8 nM (Fig. 3B). Similarly, co-incubation with sCSF1R_inh_ revealed a specific and saturable site with a K_D_ around 4.5 nM and a B_max_ indistinguishable from the other experiments (Fig. 3C). Finally, GW-2580 exhibited competitive displacement of [^3^H]5-MOP further validating the specificity of [^3^H]5-MOP (Fig. 3D).

### Inhibitor curves reveal distinct affinities of CSF1R ligands to the 5-MOP binding site

Finally, an inhibition study was done using the first human meningioma tissue sample (Fig. 4). Three compounds that are structurally distinct from 5-MOP, CPPC, sCSF1R_inh_, and GW-2580, were added to the incubation buffer to assess binding specificity and displacement properties. The resulting inhibition curves demonstrated clear differences in affinity for the [^3^H]5-MOP binding site. The calculated best-fit IC_50_ values were 66.06 nM for 5-MOP, 10.27 nM for CPPC, 1,347 nM for sCSF1R_inh_, and 122.1 nM for GW-2580.

**Figure 4.**
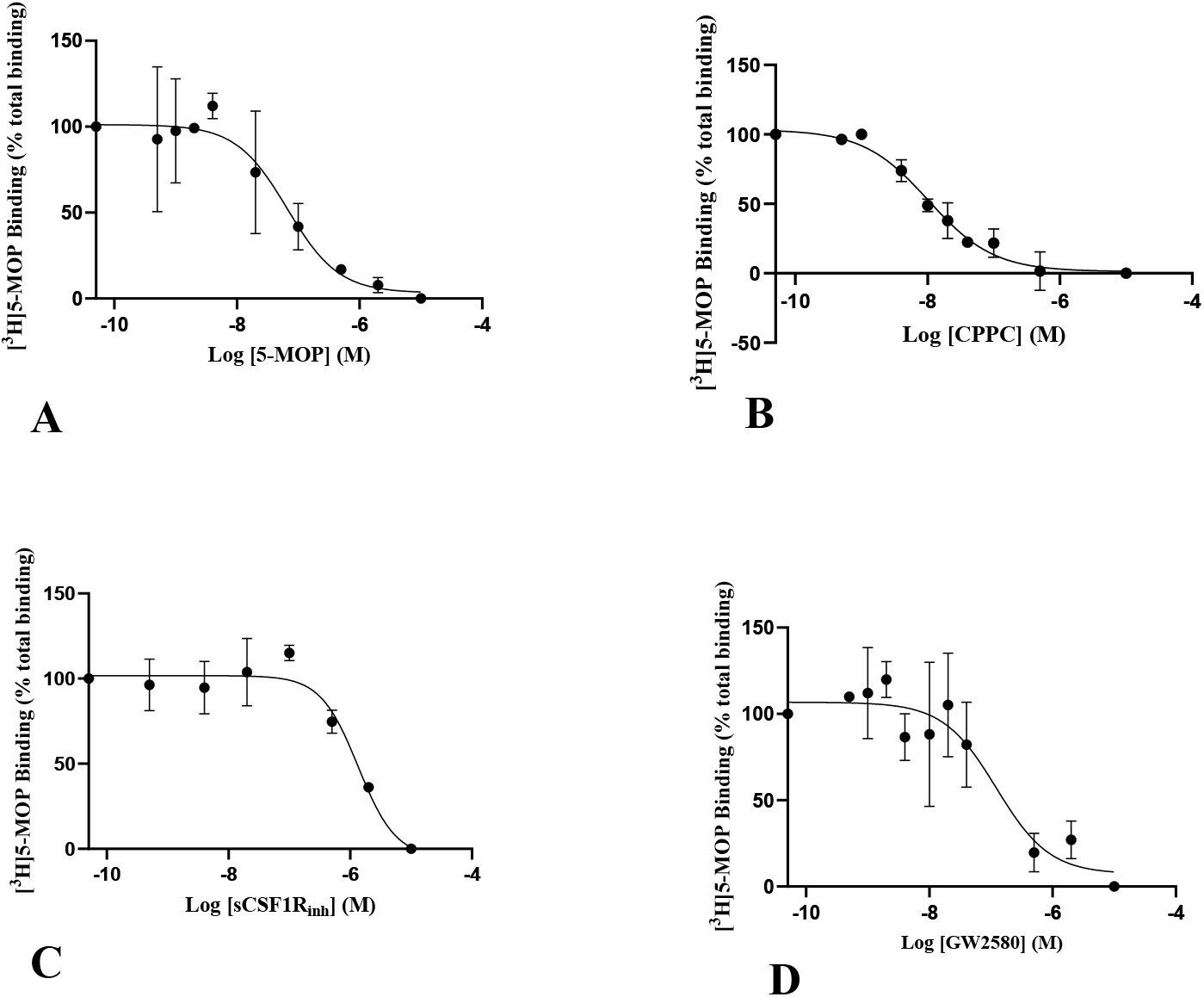
[^3^H]5-MOP is displaced by multiple CSF1R ligands in a concentration-dependent manner. Inhibitory binding curves obtained in human meningioma tissue with [^3^H]5-MOP in the presence of increasing concentrations of 5-MOP (A), CPPC (B), sCSF1R_inh_ (C), and GW-2580 (D). For all the graphs, the x-axis represents the competitor concentrations (nM), and the y-axis represents binding expressed relative to total [^3^H]5-MOP binding.

### [^3^H]CPPC binds non-CSF1R sites

The binding properties of [^3^H]CPPC were examined in the same meningioma tissue using the same approach. Qualitatively, [^3^H]CPPC showed widespread binding across the entire tissue section (Fig. 4A). Displacement with excess unlabelled CPPC reduced the signal, whereas only minimal reduction was observed in the presence of 5-MOP (Fig. 5A). The saturation binding curves generated in the absence and presence of CPPC (Fig. 5B) and 5-MOP (Fig. 5C) further indicated that [^3^H]CPPC binds to multiple non-CSF1R sites. Although [^3^H]CPPC could be displaced by the cold CPPC (Figs. 5A, 5B), the identity of the primary binding site could not be determined, as displacement by the cold ligand does not itself reveal target specificity.

**Figure 5.**
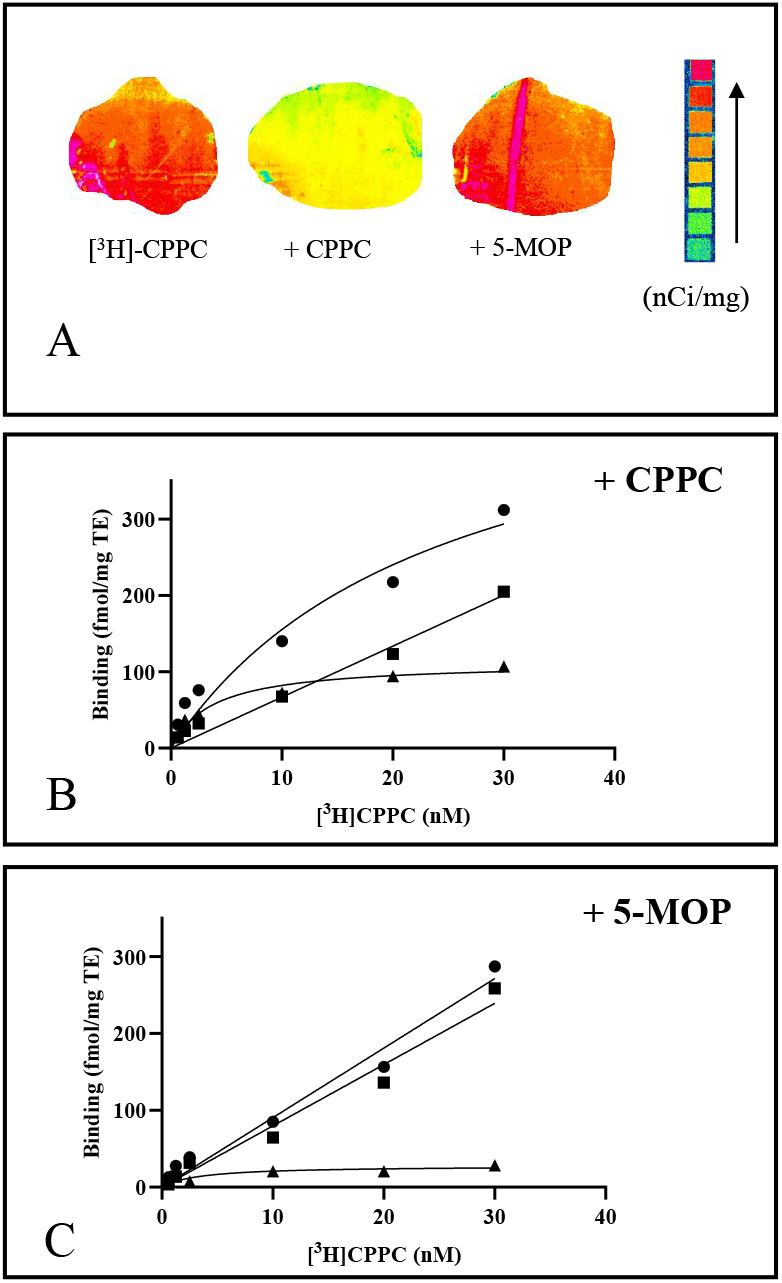
[^3^H]-CPPC binds to nonspecific sites. Representative autoradiograms (A) of human meningioma tissues incubated with [^3^H]CPPC alone or with [^3^H]CPPC in the presence of excess of CPPC or 5-MOP. Whereas CPPC displace the hot ligand, 5-MOP produced little or no displacement. Saturation binding curves generated with increasing concentrations of [^3^H]-CPPC in the presence or absence of CPPC (B) or 5-MOP (C). These experiments illustrate that displaceable binding is only observed when the unlabelled ligand matches the ratio of the radioligand itself, whereas structural distinct 5-MOP does not alter [^3^H]CPPC.

This lack of specificity was further emphasised when 5-MOP was used as a competitor where only a small fraction of total [^3^H]-CPPC was reduced in the presence of 5-MOP (Fig. 5C). The remaining signal, interpreted as putative specific binding, was consistent with the biochemical characterisation, which showed that [^3^H]-CPPC has some affinity for CSF1R, but substantial off-target binding to other kinases.

### [^3^H]5-MOP binds to activated microglia in the ischemic lesion in mice

To evaluate binding properties of [^3^H]5-MOP under pathological conditions, we also examined radioligand binding in brain sections from mice five days after an occlusion of the medial cerebral artery. The same tracer concentration of [^3^H]5-MOP was used for the human tissue experiments was applied, and binding was compared with [^3^H]CPPC and [^3^H]PBR28. Five days after the ischemic stroke, microglia become activated and infiltrate both the peri-infarct and infarct core, where necrosis is also present. Accordingly, the central zone is due to necrosis relatively low in binding. In agreement with the previous findings [^3^H]PBR28 binding was highest in the peripheral zone, and less in the central zone, probably because of intense necrosis (Zarruk *et al*., 2018) (Fig. 6). [^3^H]5-MOP binding showed a similar regional distribution although at lower intensity than for [^3^H]PBR28 (Fig. 6). In contrast, [^3^H]CPPC produced a more diffuse and widespread binding across the entire section. Relative to the surrounding tissue, the lesion appeared to have a relatively low [^3^H]CPPC signal and there was no difference in the peripheral or central zone of the lesion (Fig. 6), again suggesting binding to additional targets beyond CSF1R.

**Figure 6.**
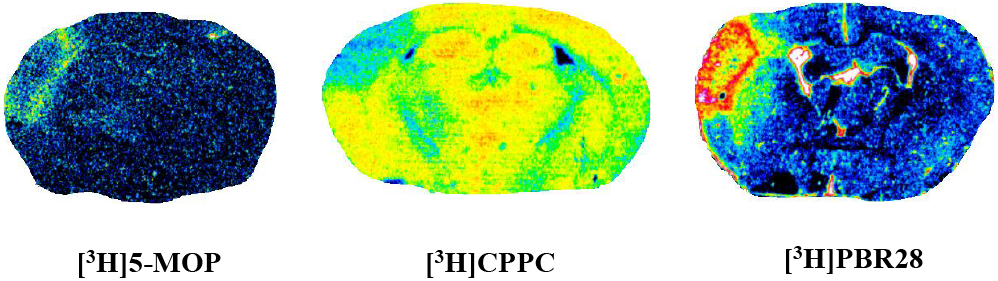
Representative autoradiograms for [^3^H]5-MOP, [^3^H]-CPPC, [^3^H]-PBR28 and in sections from an ischemic mouse model. Autoradiograms illustrate the binding patterns of [^3^H]5-MOP, [^3^H]CPPC, [^3^H]PBR28 in the ischemic mouse brain. The lesion is strongly labelled with [^3^H]-PBR28, a marker of TSPO, whereas no distinct binding is observed for [^3^H]-CPPC. The distribution of [^3^H]PBR28 labelling corresponds to the well described high density zone of activated microglia in the lesion periphery, and reduced signal in the necrotic core of the lesion. The binding pattern is the same for [^3^H]5-MOP supporting its binding to activated microglia.

## Discussion

This study demonstrated that the novel radioligand [^3^H]5-MOP selectively binds to CSF1R. Consistent with this, both the LanthaScreen kinase binding assay and broader KINOMEscan™ panel confirmed high affinity for CSF1R and excellent selectivity across a large kinase panel of 406 different kinases. The tissue binding of [^3^H]5-MOP was inhibited not only with the cold ligand itself, but also with earlier published CRF1R ligands that are structurally distinct from 5-MOP such as CPPC, sCSF1R_inh_, and GW-2580 (Altomonte *et al*., 2023;Chauveau *et al*., 2025;Das *et al*., 2026;Horti *et al*., 2019;Illig *et al*., 2008;Knight *et al*., 2021;Zhou *et al*., 2021). It would be anticipated that displacement with the cold ligand itself would reduce intensity at all sites, also those distinct from CSF1R.

The displacement with multiple structurally diverse CSF1R competitors in the autoradiography studies strongly support the specificity of [^3^H]5-MOP for CSF1R, and reveal that [^3^H]5-MOP displays no detectable off-target binding at least up to 30 nM for the tissues investigated here. This selectivity was also seen in the KINOMEscan™ panel where 5-MOP was screened against 406 kinases and showed little to minimal off-target binding.

Similar studies were conducted using CPPC, which has so far been the most widely used CRF1R radioligand including for PET studies in man (Coughlin *et al*., 2022;Horti *et al*., 2019;Mills *et al*., 2025). We also show that CPPC is binding CSF1R in the kinase screen, but it is not selective and activates a panel of other kinases at similar concentrations, some of which are expressed in the brain. This is in accordence with a previous report showing that CPPC bound to 204 out of 403 kinases tested (Knight *et al*., 2021). Furthermore, [^11^C]CPPC autoradiography revealed that the binding was only partially blocked with the CSF1R inhibitors PLX3397, BLZ945, and PLX5622, and similar binding was seen in wildtype and CSF1R knock out tissues (Horti *et al*., 2019). It is assumed that co-incubation with high concentrations of the cold ligand, CPPC, will displace binding at all sites, also those that are not CSF1R. The use of [^3^H]CPPC in inflamed tissue around an ischemic lesion show that the radioligand does not align with the region of inflammation defined by increased TSPO signal with [^3^H]PBR28 binding. Actually, [^3^H]CPPC binds broadly in the brain, which is in agreement with similar autoradiographic studies in the spleen (Knight et al. 2021). Despite earlier data demonstrating CPPC to be non-selective, the tracer has been continuously used clinically (Mills *et al*., 2025). However, it is concluded here that [^11^C]CPPC is not a reliable PET ligand for future studies.

Numerous attempts have been made to develop other PET radiotracer with high affinity, as well as good kinetics and brain-penetrating properties that selectively binds to CSF1R (Adhikari *et al*., 2024;Altomonte and Pike, 2023). The overall aim has been to identify new tools beyond TSPO that selectively determined neuroinflammatory processes in the brain, and CSF1R PET imaging is believed to support drug development by monitoring target interaction, and occupancy of novel CSF1R inhibitors sharing the same binding site (Adhikari *et al*., 2024). 5-MOP penetrates the blood-brain barrier (unpublished observations), but its application is still limited to ex vivo and in vitro studies. Another CSF1R radioligand [^11^C]GW-2580 displays good selectivity but an incomplete homologous blockade after 8 days from LPS injection and slower kinetics in mouse and monkey brains (Zhou *et al*., 2021). [^11^C]-NCGG401 was synthesized starting from BLZ945, and showed good blood brain barrier penetration and utility for PET in rodents and man (Ogata *et al*., 2025;Ogata *et al*., 2022). Autoradiography analysis of [^11^C]NCGG401 in rat brain and human postmortem hippocampal sections, did not reflect the distribution of TSPO (Ogata *et al*., 2025). Recently, JNJ-CSF1R-1 (5-cyano-N-(4-(4-(2-(fluoro)ethyl)piperazin-1-yl)-2-(piperidin-1-yl)phenyl)furan-2-carboxamide, which is a fluor analog of CPPC bound to inflamed mouse tissues (Salarian *et al*., 2025).

In this study, [^3^H]5-MOP demonstrated strong binding in brain regions with expected CSF1R upregulation following ischaemic stroke, suggesting that it could serve as a more reliable tool assessing microglia activation in disease. The comparative binding of [^3^H]5-MOP and [^3^H]PBR28 in the stroke mouse model show that the spatial distribution of [^3^H]5-MOP aligns with the penumbra region within the lesion, where microglia accumulate mostly (Zarruk *et al*., 2018). Again, [^3^H]5-MOP radioactivity was localized exactly to regions of inflammation, reinforcing its specificity for CSF1R.

Increased CSF1R expression has been demonstrated in various brain pathologies (Akiyama et al. 1994;Han et al. 2022;Olmos-Alonso et al. 2016). The validation of 5-MOP as a radioligand may also have value for drug development approaching the same target. Inhibition of CRF1R has been shown to result in removal of dysregulated microglia from the bran, reducing neuronal damage in AD animal models (Spangenberg *et al*., 2019). Further, sCSF1R_inh_ attenuated neuroinflammation after LPS in vivo, and blocked both axonal damage and behavioural impairment in an experimental autoimmune encephalomyelitis model of MS (Hagan *et al*., 2020). These findings suggest that [^3^H]5-MOP could be a more specific marker for CSF1R, particularly in conditions where CSF1R is upregulated. Radioligands have been produced to better understand and visualize microglia activation during disease progression (Altomonte *et al*., 2023;Hagan *et al*., 2020;Han *et al*., 2022;Horti *et al*., 2019;Knight *et al*., 2021;Ogata *et al*., 2022;Tanzey *et al*., 2018;Zhou *et al*., 2021). This is particular important because current targets, such as TSPO, has some limitations (Bae *et al*., 2014;Nutma *et al*., 2023). The level of binding in the meningioma probably reflecting the presence of activated microglia was sufficient for the saturation and inhibition studies here.

The inhibition of CSF1R activity leads to microglial depletion, influencing disease progression. In response to these findings, several structurally different CSF1R inhibitors have been developed to inhibit the receptor activation. Treatment with GW-2580 has been shown to significantly reduce CSF1R activation and microglia proliferation without affecting the basal levels of resting microglia (Neal *et al*., 2020).

Notably, in ischemic mouse tissues, [^3^H]5-MOP binding was confined to the lesion, with no signal detected in adjacent neuronal areas. The target cells for radioligand binding have not been identified, and it cannot be excluded that penetrating macrophages or leucocytes contribute as target cells, but they are limited in number.

We were unable to detect consistent binding in brain tissues from human and murine healthy brains. For the mice, we considered normal brain tissue to be the hemisphere contralateral to the lesion. Neither any saturation, nor displacement of the ligand could be detected. This is in accordance with the modest or any expression of CSF1R in healthy rodent brain (Michaelson *et al*., 1996).

In summary, we report that [^3^H]5-MOP is a highly specific radiotracer for CSF1R with low nanomolar affinity in both mice and humans. Given that CSF1R represents a promising drug target in neuroimmunology, 5-MOP may serve as a novel interesting tool for determining neuroinflammation and a tool for measuring target occupancy.

## List of abbreviations

5-MOP: 4-((5-methoxy-6-((5-methoxypyridin-2-yl)methoxy)pyridin-3-yl)methyl)-2-(1-methyl-1H-pyrazol-4-yl)pyrimidine
AD: Alzheimer’s disease
B_max_: Maximum binding capacity
BSA: Bovin serum albumin
CNS: Central nervous system
CPPC: 5-cyano-N-(4-(4-methylpiperazin-1 yl)-2-(piperidin-1-yl)phenyl)furan-2-carboxamide
CSF1R: Colony stimulating factor 1 receptor
GW-2580: 5-(3 Methoxy-4-((4-methoxybenzyl)oxy)benzyl)pyrimidine-2,4-dia mine
IC_50_: Half-maximal inhibitory concentration
JNJ-CSF1R-1: 5-cyano-N-(4-(4-(2-(fluoro)ethyl)piperazin-1-yl)-2-(piperidin-1-yl)phenyl)furan-2-carboxamide
KA: Kainic acid
K_d_: Constant of dissociation
MgCl_2_: Magnesium chloride
MS: Multiple sclerosis
mTLE: Medial temporal lobe epilepsy
PBR28: N-(2-methoxybenzyl)-N-(4-phenoxypyridn-3-yl)acetamide
PD: Parkinson’s disease
PET: Positron emission tomography
pMCAo: Permanent medial cerebral artery occlusion
PK11195: 1-(2-chlorophenyl)-N-methyl-N-(1-methylpropyl)-3-isoquinoline carboxamide
sCSF1R_inh_: (S)-4-(3-((2-(6-methoxypyridin-3-yl)-2,3-dihydrobenzo[b]dioxin-6-yl)methyl)-3H-imidazo[4,5-b]pyridin-6-yl)-2-methylbut-3-yn-2-amine
TE: Tissue equivalent
TR-FRET: Time-resolved fluorescence energy transfer
TSPO: Translocator protein, 18kDa

## Author contribution

Conceptualization: BBA, JFB, BF, JDM;Methodology: CI, BAP, BBA, MH, TJ, KL, AL, Formal analysis: CI, BAP, MH, TJ;Resources: JFB and JDM.

JDM wrote the original draft and all authors commented and approved the final draft.

